# SARS-CoV-2 Spike receptor-binding domain with a G485R mutation in complex with human ACE2

**DOI:** 10.1101/2021.03.16.434488

**Authors:** Claire M. Weekley, Damian F. J. Purcell, Michael W. Parker

## Abstract

Since SARS-CoV-2 emerged in 2019, genomic sequencing has identified mutations in the viral RNA including in the receptor-binding domain of the Spike protein. Structural characterisation of the Spike carrying point mutations aids in our understanding of how these mutations impact binding of the protein to its human receptor, ACE2, and to therapeutic antibodies. The Spike G485R mutation has been observed in multiple isolates of the virus and mutation of the adjacent residue E484 to lysine is known to contribute to antigenic escape. Here, we have crystallised the SARS-CoV-2 Spike receptor-binding domain with a G485R mutation in complex with human ACE2. The crystal structure shows that while the G485 residue does not have a direct interaction with ACE2, its mutation to arginine affects the structure of the loop made by residues 480-488 in the receptor-binding motif, disrupting the interactions of neighbouring residues with ACE2 and with potential implications for antigenic escape from vaccines, antibodies and other biologics directed against SARS-CoV-2 Spike.

## Introduction

Following its emergence in 2019, the evolution of SARS-CoV-2 has been monitored by genomic sequencing. Many point mutations have been identified in the receptor-binding domain of the SARS-CoV-2 Spike glycoprotein (Spike RBD) that binds angiotensin-converting enzyme 2 (ACE2) on the surface of human cells as part of the process of cellular entry and viral infection. Some of these mutations alone and in combination with other mutations in the Spike protein have been associated with increased transmissibility or infectivity of the virus that causes COVID-19 (Wu *et al.*, 2021). Mutations in the receptor-binding domain of the Spike protein may also enable antibody escape and reduce the efficacy of vaccine-elicited immunity (Williams and Burgers, 2021).

The G485 residue is not directly involved in interactions with ACE2 residues, but it is found in the B1’/B2’ loop region (residues 455-491) of the receptor-binding motif that is notable for its high sequence variation between the SARS-CoV and SARS-CoV-2 Spike RBD. Wang *et al.* (2020) identified 115 van der Waals contacts to ACE2 in this SARS-CoV-2 B1’/B2’ loop region, in addition to three potential hydrogen bonds. G485 has been identified amongst antibody binding epitopes in the RBD (Kreye *et al.*, 2020; Greaney *et al.*, 2021). Its neighbouring residue E484 is of great interest to the scientific community with the mutation E484K contributing to antigenic escape (Greaney *et al.*, 2021; Liu *et al.*, 2021). The mutation of residue 485 from glycine to arginine has been observed in SARS-CoV-2 sequenced from isolates in China in February 2020 (Nguyen *et al.*, 2021) and in isolates associated with an outbreak among abattoir workers in Australia documented in the GISAID database (unpublished data; *e.g.* GISAID name: hCoV-19/Australia/VIC1588/2020; Accession ID: EPI_ISL_456508, Victorian Infectious Diseases Reference Laboratory).

Studies of the effect of the G485R mutation on Spike function are limited. In a deep mutational scanning study of the receptor-binding domain, the G485R mutation cause only a slight reduction in affinity of the Spike RBD for ACE2 (Starr *et al.*, 2020). However, the G485R mutation was found to reduce viral neutralisation of some convalescent plasma up to 5-fold (Greaney *et al.*, 2021).

The location of G485 in a loop region critical for interaction with ACE2 and the reported antigenic effect of the G485R mutation prompted us to determine the structure of the G485R Spike RBD in complex with ACE2.

## Materials and Methods

### Plasmids

Residues 1-617 of human ACE2 ectodomain (UniProtKB Q9BYF1C) and SARS-CoV-2 spike residues 319-530 (GenBank QHD43416.1) with the G485R mutation, were each synthesised with a N-terminal honeybee melittin signal peptide and a C-terminal TEV protease cleavage site and His_6_-tag by GenScript (Hong Kong) and subcloned into the pFastBac1 vector.

### Protein expression and purification

ACE2 and Spike RBD protein were expressed in Sf21 cells using the Bac-to-Bac system. Cells at a density of 2 × 10^6^ cells/mL were infected with 1:1000 V2 viral stock. Supernatants were collected 44-48 h (ACE2) and 68-72 h (Spike RBD) post-infection, centrifuged and loaded onto a HisTrap Excel column. The protein was eluted with 50 mM Tris pH 7.5 and 150 mM NaCl along a linear gradient to 500 mM imidazole. Spike RBD was purified by size exclusion chromatography on a Superdex 200 10/300 Increase GL in 20 mM Tris pH 7.5 and 150 mM NaCl. The ACE2 His_6_-tag was cleaved using TEV protease and removed by Ni-NTA chromatography. To form the Spike RBD-ACE2 complex, an excess of G485R Spike RBD was added to ACE2, incubated for 30 minutes at room temperature and the purified complex was isolated by size exclusion chromatography as described above.

### Crystallisation and structure determination

The G485R Spike RBD-ACE2 complex was concentrated to 5 mg/mL through a 10 kDa molecular weight cut-off filter and added in a 1:1 ratio with crystallisation buffer in a sitting drop plate. Several hits were obtained using the MCSG-1 (Anatrace) screen and crystallisation was optimised around MCSG condition A1. The crystal used to determine the structure was obtained in 0.1 M HEPES pH 7.0 and 16% w/v PEG 8000 at 293 K. The crystal was transferred to cryobuffer consisting of 20% ethylene glycol added to the mother liquor and flash frozen. Data were collected at 100 K at the Australian Synchrotron’s MX2 beamline using the ACRF Dectris EIGER 16M detector (Aragão *et al.*, 2018; Phillips *et al.*, 2002). Data were integrated using XDS and scaled with AIMLESS from the CCP4 program suite (Winn *et al.*, 2011). Molecular replacement was performed using Phaser from the Phenix program suite with the SARS-CoV-2 Spike RBD-ACE2 structure (PDB ID: 6LZG) used as a search model. Alternate rounds of model building and refinement were performed using Coot (Emsley *et al.*, 2010), REFMAC5 (Murshudov *et al.*, 2011) and phenix.refine (Afonine *et al.*, 2012). The refined structure has been deposited in the Protein Databank with PDB ID: 7LO4. The diffraction data and refinement statistics are summarised in Table 1.

**Table 1.**
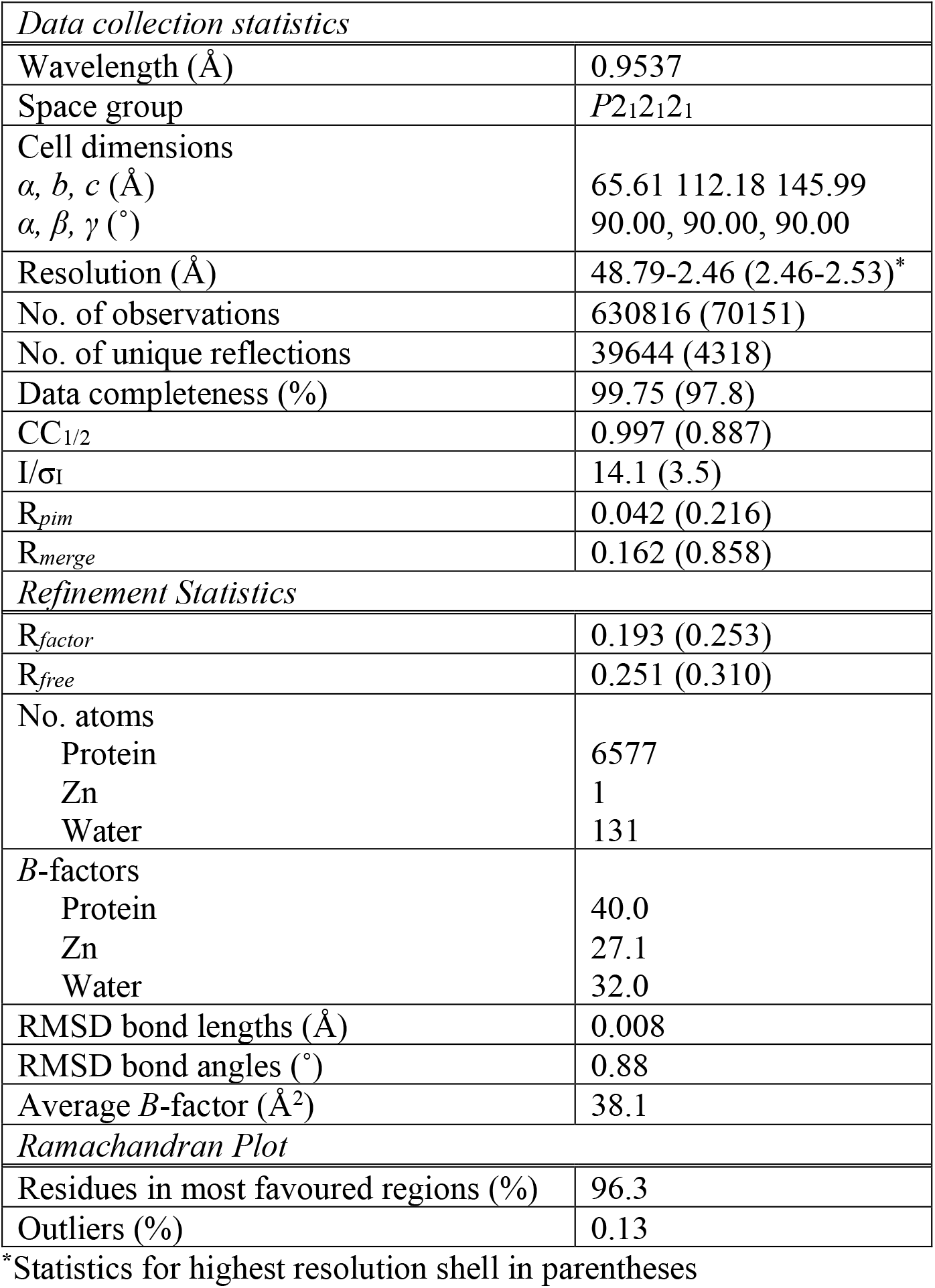
Crystallographic data collection and refinement statistics (PDB ID: 7LO4)

## Results and Discussion

Attempts to crystallise the G485R Spike RBD-ACE2 complex under conditions previously reported for SARS-CoV Spike RBD (Li *et al.*, 2005) and SARS-CoV-2 Spike RBD (Wang *et al.*, 2020; Lan *et al.*, 2020) resulted in either no crystals or non-diffracting crystals. A crystal diffracting to 2.5 Å resolution was ultimately obtained from 0.1 M HEPES pH 7.0 and 16% w/v PEG 8000. The complex crystallised in the *P*2_1_2_1_2_1_ space group (compared to the *P*4_1_2_1_2 space group of the wild-type complexes) with one complex in the asymmetric unit. We determined the structure using standard molecular replacement and refinement methods.

The electron density maps supported that Spike RBD is glycosylated at N343 as expected (Watanabe *et al.*, 2020) and observed previously in crystal structures (Wang *et al.*, 2020). In the ACE2 molecule, a single zinc ion is coordinated by residues H374, H378 and E402. There is sufficient density to model glycans at residues N53, N90, N103 and N546 – four of the six known N-glycosylation sites in the ACE2 construct (Shajahan *et al.*, 2020) (Figure 1a).

**Figure 1.**
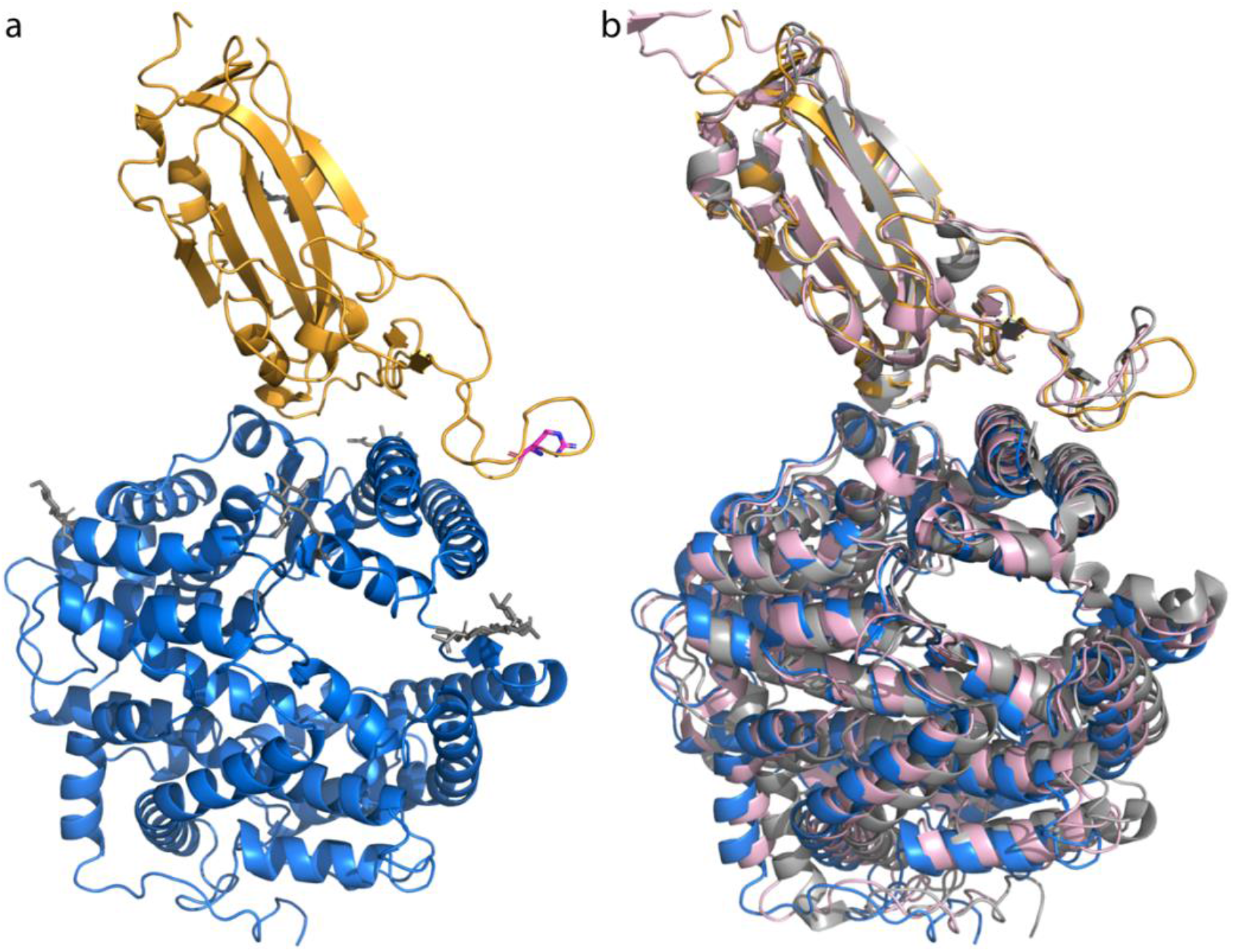
Crystal structure of G485R Spike RBD bound to ACE2. (a) Ribbon picture of the complex. The G485R Spike RBD and ACE2 are coloured orange and blue, respectively. Residue R485 is highlighted in pink. Glycosylated residues are shown in dark grey sticks. (b) Comparison to the previously reported wild-type crystal structure (PDB ID: 6M0J) and cryo-electron microscopy structure (PDB ID: 7KJ2). The complexes have been superimposed using the Spike RBD domain. The 6M0J structure is coloured light grey and the 7KJ2 structure is in pink.

The mutated residue G485 is part of the B1’/B2’ loop region (residues 455-491) that makes multiple contacts with ACE2. The smaller loop within this region (residues 480-488), created by the disulfide bond between C480 and C488, is rotated in the mutant structure relative to the wild-type Spike-RBD ACE2 crystal structure and a cryo-electron microscopy structure of the full-length Spike trimer with one ACE2 bound (Figure 1b) (Xiao *et al.*, 2021). The root mean square deviation between the wild-type (PDB ID: 6M0J) and G485R structures is 2.2 Å over the 9 C _α_ atoms in residues 480-488 compared to 1.63 Å over all 1536 C _α_ atoms. Due to the rotation, only one of the two hydrogen bonds between the G485 and C488 backbone is retained with the mutation to R485 and the C488 ϕ and ψ angles change from −136° and 102°, respectively, in the wild-type structure to −119° and 148° in the mutant structure. Notably, the orientations of residues E484 and F486 are shifted away from the ACE2 interface (Figure 2). In the wild-type structure F486 is engaged in hydrophobic interactions with L79, M82 and M83 of ACE2 and E484 forms a salt bridge with the K31 residue of ACE2 (Wang *et al.*, 2020). In the G485R mutant structure, these interactions are disrupted (Figure 2b and 2c) – residue 484 in the G485R structure is rotated ~160° compared to the wild-type structure – and there are no additional interactions between the B1’/B2’ loop residues and ACE2 to compensate (Table 2).

**Table 2.** List of polar and non-polar contacts between the Spike RBD and ACE2.

| Spike residue | ACE2 residue | Number of contacts |  |  |
| --- | --- | --- | --- | --- |
|  |  | G485R (7LO4) | WT (6M0J) | WT (6LZG) |
| K417 | D30 | 3(1)<sup>a</sup> | 4(1) | 3(1) |
| G446 | Q42 | 1 | 1(1) | 1 |
| Y449 | D38 | 6(1) | 6(1) | 5(2) |
|  | Q42 | 1(1) | 1(1) | 1(1) |
| Y453 | H34 | 5(1) | 3 | 3 |
| **L455** | H34 | - | 2 | 4 |
| **F456** | T27 | 5 | 5 | 5 |
|  | D30 | 1 | 1 | 1 |
|  | K31 | - | - | 1 |
| **A475** | S19 | 3(1) | - | 3(1) |
|  | Q24 | 1 | 1 | 1 |
|  | T27 | 1 | 0 | 1 |
| **G476** | S19 | 1 | - | 2 |
|  | Q24 | 3 | - | - |
| **E484** | K31 | - | 1<sup>b</sup> | 1<sup>c</sup> |
| **F486** | L79 | - | 1 | - |
|  | M82 | - | 4 | 4 |
|  | Y83 | - | 6 | 7 |
| **N487** | Q24 | 3(1) | 4(1) | 7(1) |
|  | Y83 | 4(1) | 5(1) | 4(1) |
| **Y489** | T27 | 2 | 1 | 2 |
|  | F28 | 3 | 3 | 4 |
|  | K31 | 1 | 1 | - |
|  | Y83 | 1 | 1 | 1 |
| **F490** | K31 | - | - | 1 |
| Q493 | K31 | 5(1) | 3 | - |
|  | H34 | 4 | 6 | 4 |
|  | Q35 | 4(1) | 4(1) | 4(1) |
| S494 | H34 | 2(1) | - | - |
| G496 | D38 | - | - | 1 |
|  | K353 | 3(1) | 4(1) | 1(1) |
| Q498 | L45 | - | - | 2 |
|  | Q38 | 4(1) | - | 1 |
|  | Y41 | 6 | 6 | 5 |
|  | Q42 | 3(0) | 3(1) | 9(1) |
|  | K353 | 5(1) | - | - |
| T500 | Y41 | 6(1) | 5(1) | 6(1) |
|  | N330 | 1 | 4 | 3 |
|  | D355 | 6 | 5 | 2 |
|  | R357 | 3 | 3 | 3 |
| N501 | Y41 | 5(1) | 4(1) | 4(1) |
|  | K353 | 6 | 3 | 5 |
| G502 | K353 | 3(1) | 3(1) | 3(1) |
|  | G354 | 6 | 4 | 5 |
|  | D355 | 1 | - | - |
| Y505 | E37 | 8(1) | 3(1) | 3 |
|  | K353 | 15 | 14 | 15 |
|  | G354 | 2 | 2 | 2 |
|  | R393 | 1 | 1 | - |

<sup>a</sup>Number of non-polar contacts (4 Å cut-off) and, in parentheses, number of potential hydrogen bonds (3.5 Å cut-off). Residues in bold are in the B1'/B2' loop region. <sup>b</sup>Corresponding interaction in 6LZG identified as a salt bridge by Wang *et al.* (2020); interaction measured at 4.4 Å. <sup>c</sup>Interaction measured at 4.2 Å; reported as a salt bridge by Wang *et al.* (2020).

**Figure 2.**
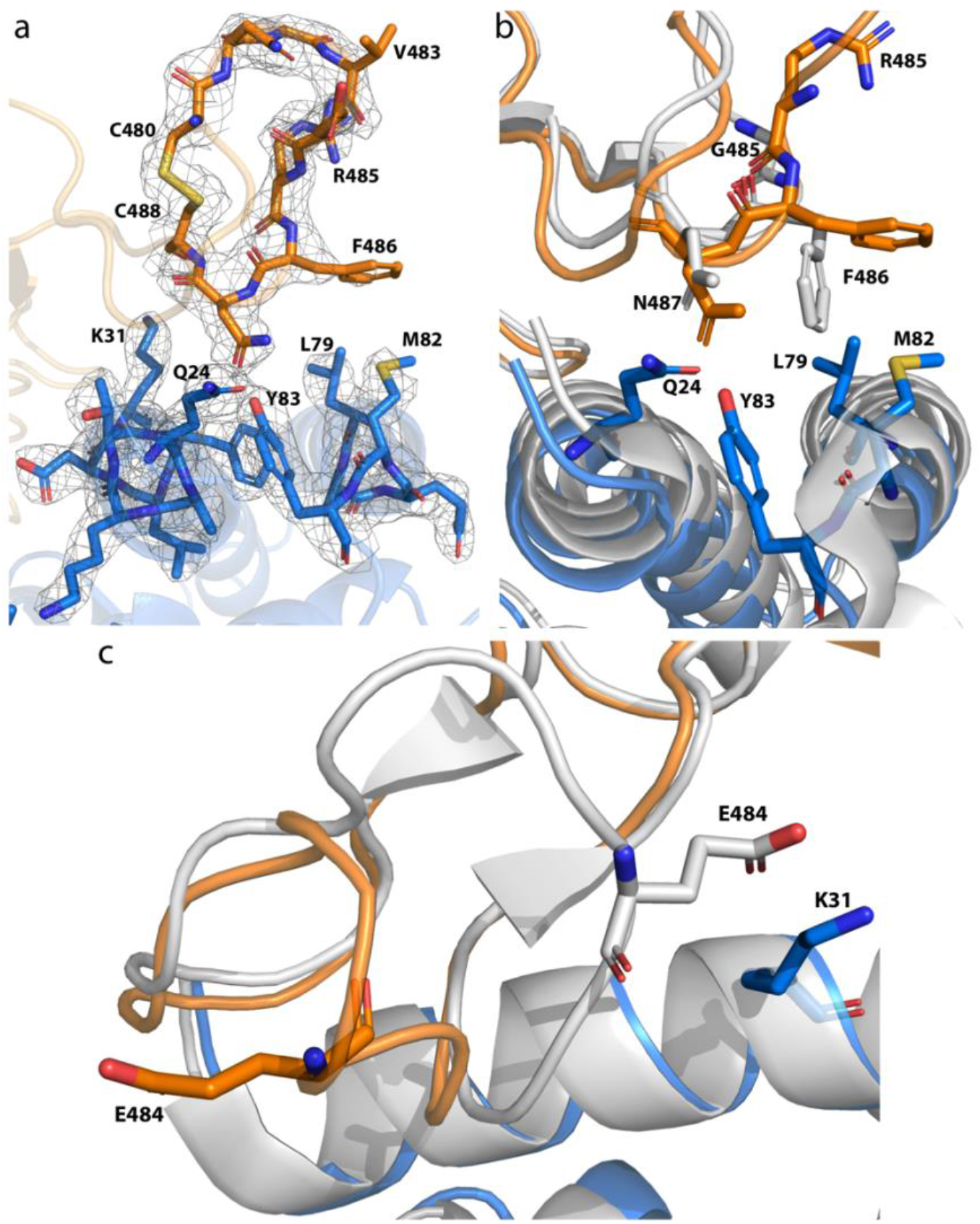
Details of B1’/B2’ loop residues in the Spike RBD receptor-binding domain and their interactions with ACE2. (a) The electron densities of G485R Spike RBD (orange) residues 480-488 and ACE2 (blue) residues 24-31 and 79-83 are shown in grey mesh with the 2*F*_*o*_–*F*_*c*_ map contoured at 1σ. Comparison of the G485R Spike RBD-ACE2 structure to the wild-type structure in grey (PDB ID: 6M0J) reveals the shift in the orientations of Spike RBD residues (b) F486 and (c) E484 away from ACE2 residues.

The number and type of crystal lattice contacts also differs between the wild-type and mutant. In the wild-type structure the Spike RBD N481, G482, E484 and F490 residues form crystal contacts with ACE2 residues N250, N251 and N601 in the neighbouring molecule (Table 3). In the mutant structure, an entirely different set of crystal contacts are observed, including extensive interactions between R485 and residues D597, Q598, K600 and N601 that may stabilise R485 and the C480-C488 loop in its observed orientation. Therefore, we cannot rule out the contribution of crystal lattice artifacts to the structural changes observed with the G485R mutation.

**Table 3.** List of contacts between the Spike RBD residues 480-490 and the neighbouring molecules in the crystal lattice.

| Spike residue | Residue from neighbouring molecule | Number of contacts |  |  |
| --- | --- | --- | --- | --- |
|  |  | G485R (7LO4) | WT (6M0J) | WT (6LZG) |
| N481 | N601 | - | 0/1<sup>a</sup> | 0/2 |
| G482 | N250 | - | 1/2 | 1/1 |
| E484 | N250 | - | 0/1 | 1/1 |
| R485 | D597 | 2/6 | - | - |
|  | Q598 | 1/4 |  |  |
|  | K600 | 0/1 |  |  |
|  | N601 | 4/7 |  |  |
| C488 | N601 | 2/3 | - | - |
| F490 | N250 | - | 1/8 | 0/7 |
|  | N251 | - | 0/2 | 0/1 |
|  | I259 | 0/2 | - | - |
|  | N601 | 0/2 | - | - |
|  | N602 | 0/3 | - | - |
|  | N603 | 0/5 | - | - |

<sup>a</sup>Contacts $\leq 3.5$ Å/Contacts between 3.5 and 4 Å.

Despite the structural evidence that the G485R mutation reorients residues 484 and 486 away from ACE2 and reduces the number of interactions between the Spike RBD and ACE2, biophysical studies indicate that this mutation is not sufficient to appreciably reduce the affinity of the Spike RBD for ACE2 (Starr *et al.*, 2020). Thus it would not be expected to reduce the transmissibility of a virus carrying this mutation. Indeed, a G485R mutation arising in a high risk transmission setting in Australia was documented in 37 recipients (unpublished data).

Mutations of residues in the B1’/B2’ loop region of Spike RBD have been shown to be involved in antigenic escape from neutralising antibodies. Mutations at E484 and/or F486 arose as escape substitutions multiple times when a SARS-CoV-2 virus was tested against a panel of neutralising monoclonal antibodies against the SARS-CoV-2 RBD (Liu *et al.*, 2021). The structure of the Spike-RBD in complex with two monoclonal antibodies in an investigational antibody cocktail have recently been published (Dong *et al.*, 2021). These structures reveal that Spike RBD residues 485-487 are surrounded by a hydrophobic pocket formed by the monoclonal antibody COV2-2196, revealing the importance of these residues in antibody binding. Evidence has also emerged that G485R also plays a role in antigenic escape from neutralising antibodies isolated from SARS-CoV-2 convalescent patients, with mutations at F486 also having an impact (Greaney *et al.*, 2021). This finding correlates with our structural data which reveals significant conformational rearrangements in the loop, including disruptions of interactions involving E484 and F486 with the G485R mutation.

## Supplementary Information

The sequences of the protein constructs and evidence of G485R Spike RBD-ACE2 complex formation from size exclusion chromatography are included in the supplementary file.

## Acknowledgements

This research was undertaken in part using the MX2 beamline at the Australian Synchrotron, part of ANSTO, and made use of the Australian Cancer Research Foundation (ACRF) detector. We gratefully acknowledge funding from the Victorian Government under the COVID-19 Victorian Consortium. We thank Dr Michael Gorman, Dr Larissa Doughty and Dr Craig Morton for helpful discussions and advice.

## Notes

### Competing Interest Statement

The authors have declared no competing interest.

