## Supplementary Information for "SARS-CoV-2 Spike receptor-binding domain with a G485R mutation in complex with human ACE2"

The protein constructs used in this work are as follows:

G485R SARS-CoV-2 Spike RBD Construct

```
MKFLVNVALV FMVVYISYIY ARVQPTESIV RFPNITNLCP FGEVFNATRF ASVYAWNRKR
ISNCVADYSV LYNSASFSTF KCYGVSPTKL NDLCFTNVYA DSFVIRGDEV RQIAPGQTGK
IADYNYKLPD DFTGCVIAWN SNNLDSKVGG NYNYLYRLFR KSNLKPFERD ISTEIYQAGS
TPCNGVERFN CYFPLQSYGF QPTNGVGYQP YRVVVLSFEL LHAPATVCGP KKSENLYFQG
GSHHHHHH
```

Human ACE2 Construct

```
MKFLVNVALV FMVVYISYIY ASSSSWLLLS LVAVTAAQST IEEQAKTFLD KFNHEAEDLF
YQSSLASWNY NTNITEENVQ NMNNAGDKWS AFLKEQSTLA QMYPLQEIQN LTVKLQLQAL
QQNGSSVLSE DKSKRLNTIL NTMSTIYSTG KVCNPDNPQE CLLLEPGLNE IMANSLDYNE
RLWAWESWRS EVGKQLRPLY EEYVVLKNEM ARANHYEDYG DYWRGDYEVN GVDGYDYSRG
QLIEDVEHTF EEIKPLYEHL HAYVRAKLMN AYPSYISPIG CLPAHLLGDM WGRFWTNLYS
LTVPFGQKPN IDVTDAMVDQ AWDAQRIFKE AEKFFVSVGL PNMTQGFWEN SMLTDPGNVQ
KAVCHPTAWD LGKGDFRILM CTKVTMDDFL TAHHEMGHIQ YDMAYAAQPF LLRNGANEGF
HEAVGEIMSL SAATPKHLKS IGLLSPDFQE DNETEINFLL KQALTIVGTL PFTYMLEKWR
WMVFKGEIPK DQWMKKWWEM KREIVGVVEP VPHDETYCDP ASLFHVSNDY SFIRYYTRTL
YQFQFQEALC QAAKHEGPLH KCDISNSTEA GQKLFNMLRL GKSEPWTLAL ENVVGAKNMN
VRPLLNYFEP LFTWLKDQNK NSFVGWSTDW SPYADQSENL YFQGGSHHHH HH
```

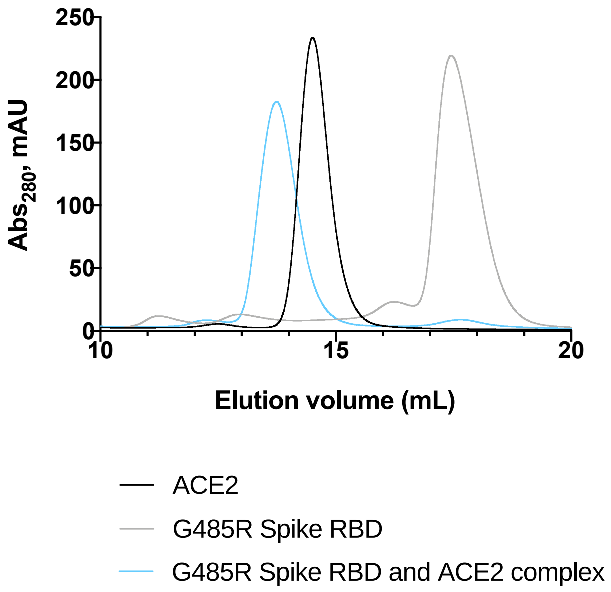


**Supplementary Figure 1.** The formation of the G485R Spike RBD-ACE2 complex was confirmed by size exclusion chromatography.
